# Biodegradable and Biocompatible Graphene-based Scaffolds for Functional Neural Tissue Engineering: A Strategy Approach Using Dental Pulp Stem Cells and Biomaterials

**DOI:** 10.1101/2021.01.12.426431

**Authors:** Negar Mansouri, Said Al-Sarawi, Dusan Losic, Jagan Mazumdar, Jillian Clark, Stan Gronthos, Ryan O’Hare Doig

## Abstract

Neural tissue engineering aims to restore function of nervous system tissues using biocompatible cell-seeded scaffolds. Graphene-based scaffolds combined with stem cells deserve special attention to enhance tissue regeneration in a controlled manner. However, it is believed that minor changes in scaffold biomaterial composition, internal porous structure, and physicochemical properties can impact cellular growth and adhesion. The current work aims to investigate *in vitro* biological effects of 3D graphene oxide (GO)/sodium alginate (GOSA) and reduced GOSA (RGOSA) scaffolds on dental pulp stem cells (DPSCs) in terms of cell viability and cytotoxicity. Herein, the effects of the 3D scaffolds, coating conditions, and serum supplementation on DPSCs functions are explored extensively. Biodegradation analysis revealed that addition of GO enhanced the degradation rate of composite scaffolds. Compared to the 2D surface, the cell viability of 3D scaffolds was higher (p <0.0001), highlighting the optimal initial cell adhesion to the scaffold surface and cell migration through pores. Moreover, the cytotoxicity study indicated that the incorporation of graphene supported higher DPSCs viability. It is also shown that when the mean pore size of scaffold increases, DPSCs activity decreases. In terms of coating conditions, poly-l-lysine (PLL) was the most robust coating reagent that improved cell-scaffold adherence and DPSCs metabolism activity. The cytotoxicity of GO-based scaffolds showed that DPSCs can be seeded in serum-free media without cytotoxic effects. This is critical for human translation as cellular transplants are typically serum-free. These findings suggest that proposed 3D GO-based scaffolds have favourable effects on the biological responses of DPSCs.

## Introduction

The biological, neurochemical and anatomical complexity of the human nervous system challenges attempts to achieve neuronal repair or regeneration after disease or traumatic injury. The over-arching goal is to restore the functional properties of the nervous system, compensate or substitute for tissue defects, and restore neural transmission [1]. In relation to these goals, it is of considerable interest that bioengineered scaffolds can induce topographical, chemical and biological cues that effectively stimulate nerve regeneration [2]. However, there is consensus that natural microenvironments and their*in vivo* physiological equivalent conditions cannot be fully represented using two-dimensional (2D) cell cultures. Three-dimensional (3D) cell models are a more reliable representation of the physiological environment of living tissue which suitably replicate the *in vivo*native matrix and mimic the biophysical properties of remnant tissue [3, 4].

The 3D cell culture products in tissue engineering development include porous scaffolds, scaffold-free constructs, (self-assembling) hydrogels, and microchips. Among these, pre-fabricated porous scaffolds are recognized as the most promising platform to advance cell therapy and drug discovery [5]. These scaffolds are used as a supportive matrix, replicating the extracellular matrix of the central nervous system (CNS). *In vitro* proof-of-principle work shows that 3D interstices are conducive to cell attachment, migration, and infiltration [6]. A tissue-engineered scaffold with suitable biocompatibility, biodegradability and interconnected porosity is favourable in neural tissue engineering (NTE) applications [7].

The literature presents multiple examples of polymer-based materials of relevance to the fabrication of 3D scaffolds. Of these, the non-toxic, biodegradable and biocompatible properties of alginate, a natural biopolymer, have been extensively investigated for neural applications [8]. The 3D alginate-based scaffolds of Ansari et al. (2017) demonstrated efficacy to sustainably release neurotrophic factors and enhance the proliferation and neurogenic differentiation of encapsulated mesenchymal stem cells (MSCs) *in vitro* [9]. Another *in vitro* study by Wang et al. (2017) demonstrated the aptitude of hybrid scaffolds composed of chitosan and alginate to promote olfactory ensheathing and neural stem cell (NSC) viability [10]. Similarly, an*in vivo* study by Prang et al. demonstrated the compatibility of alginate-based scaffolds loaded with NSCs to dampen inflammatory response in experimental spinal cord injury necessary to encourage axonal regrowth [11]. Despite promising results, *in vitro* and*in vivo*, alginate-based constructs suffer from weak mechanical strength, high degradation rate and electrical insulation at biological frequencies, which might be attuned through combination with other biomaterials [12].

Recently, graphene-based scaffolds have attracted intense interest for their use in NTE due to their unique properties including large surface area, excellent electrical conductivity, suitable biocompatibility, chemical stability, and mechanical properties [13]. Importantly, the high electrical conductivity of graphene provides a great electrical coupling between regenerating nerve cells which is conducive to the regeneration of excitable tissues [14]. Two graphene derivatives, namely graphene oxide (GO) and reduced graphene oxide (RGO) are endowed with unique physicochemical properties of interest to functional NTE [15, 16]. Serrano et al. [17] showed that GO scaffolds improve the differentiation of NSCs into mature neurons replete with axons, dendrites and synapses and supportive glial cells. In addition, researchers observed that biological properties and cytotoxicity of graphene-based composites could be enhanced with respect to cell type, the interaction between graphene and the matrix, graphene concentration and composite production method [18]. These data suggest that graphene-based composites may warrant closer investigation.

Details on the physicochemical characterization of the GO powder used as well as of the obtained rGO scaffolds were published elsewhere. [35]

Numerous studies showed that coating reagents such as poly-l-lysine (PLL) and laminin (LAM) on the surface of the scaffolds induce signals regulating cell responses, adhesion and growth [19]. However, it is interesting to point out that various coating reagents have different impacts on cellular behaviour according to scaffold biomaterial and cell type [20]. Culture medium is another consideration, with more information required about the interaction between serum, protein corona, and scaffold properties [21-23]. These data suggest a requirement for well-designed laboratory protocols, taking into account these considerations.

We have previously reported the mechanical, electrical and physical properties of engineered 3D composite scaffolds consisting of GO and sodium alginate (SA) (GOSA) [24]. Our study revealed that GOSA composite porous scaffolds combine the known advantages of alginate (including non-toxicity, biocompatibility, biodegradability) and graphene (including hydrophilicity, excellent mechanical strength, suitable biocompatibility, good electrical conductivity). The next step was to realize that the incorporation of GO into SA to produce a composite scaffold introduces excellent chemical properties, mechanical strength and electrical conductivity, which perhaps can be harnessed to exploit the CNS physiology. In our previous work, it was shown that GOSA and RGOSA scaffolds with 0.5 and 1 wt. (%) concentrations and mean pore sizes of 147.4 µm (GOSA0.5), 142.5 µm (GOSA1), 116.0 µm (RGOSA0.5), and 114.7 µm (RGOSA1) can accommodate stem cells, as an effective and promising cell source in regenerative therapies, in culture. Therefore, it will be important to establish cellular viability and cytotoxicity data based upon potential mechanisms of graphene-based material incorporation into the scaffold.

Various studies have demonstrated the significant neurotrophic expression and secretion of DPSC encompassing nerve growth factor (NGF), brain-derived neurotrophic factor (BDNF), neurotrophin-3 (NT-3), glial cell-line derived neurotrophic factor (GDNF), vascular endothelial growth factor (VEGF) and platelet-derived neurotrophic factor (PDGF) [12, 25–27].

Neural crest-derived human dental pulp stem cells (hDPSCs), are a rich source of mesenchymal stem cells (MSCs), with the ability for high proliferation and multi-lineage differentiation capacity [25]. These cells can be easily harvested from human exfoliated deciduous teeth, permanent and primary teeth, and supernumerary teeth [26]. It has also been shown that proliferation and cell number of DPSCs are greater than bone marrow-derived MSC [27]. Studies showed that hDPSCs can differentiate into neuron-like cells and form functionally active neurons, under the direction of appropriate environmental cues [28, 29]. DPSCs also appear to induce axonal guidance via stromal-derived factor-1 (SDF-1) secretion, encouraging further exploration. Another study by Nosrat et al. [30] showed that DPSCs express repertoire of neurotrophic factor and stimulate neurogenesis and angiogenesis [31, 32]. Similarly, implantation of hDPSCs has been shown to significantly improve forelimb sensorimotor function in cerebral ischemia rodent model [33]. For all these reasons, hDPSCs represent a candidate stem cell population for *in vitro* investigation of neural tissue repair.

Following the fabrication of composite graphene-based scaffolds, in this paper, we present our investigations into the influence of graphene incorporation, coating conditions, and DPSC donor type on the viability and functions of DPSCs. The viability and cytotoxicity of DPSC-loaded GOSA and RGOSA scaffolds have been assessed using the Alamar blue (AB) and lactate dehydrogenase (LDH) activity assays. Furthermore, defined serum-free media has been developed for the culture of DPSCs on the fabricated scaffolds to overcome the problematic issues of using fetal bovine serum (FBS) and make efficient clinical translations of stem cell-based approaches.

In previous work, we engineered vascular networks on biocompatible and biodegradable poly(?-lactic acid)(PLLA)/polylacticglycolic acid (PLGA) sca?olds, using a coculture of endothelial cells and support cells

## Experimental Section

### 2.1. Materials

Sodium alginate and Calcium chloride dried were purchased from Chem-Supply. Alpha-MEM (Gibco, Life Technologies, Australia, Cat. No. 12561056) was supplemented with fetal bovine serum (FBS, USA origin, Life Technologies, Australia), penicillin /streptomycin (Gibco), L-ascorbic acid-2-phosphate (Sigma-Aldrich), and 2mM L-Glutamine (Life Technologies, Australia). Natural mouse Laminin (Gibco-Life Technologies) and 0.01% Poly-l-lysine (PLL) solution (Sigma) were used as coating reagents. Trypsin-EDTA was supplied by Gibco Life Technologies, Australia. Clear-bottom 24-well and 96-well plates (Costar®, Corning, NY, USA) were used throughout the study. The cytotoxicity assay was performed following the manufacturer’s protocol using LDH kit from Promega Corporation (WI, USA). The AlamarBlue™ cell viability reagent was supplied by Invitrogen–ThermoFisher Scientific, USA.

### 2.2. GOSA/RGOSA Scaffold Fabrication and Preparation

The graphene-based composite scaffolds were fabricated by a technique involving solution mixing, freeze-drying, crosslinking, and bio-reduction as described previously [24]. Briefly, 2 wt. % sodium alginate powder was dissolved in deionized water until a transparent solution is obtained. Then, GO (synthesized by modified hummers’ method as described earlier [34]) was added to the solution to get a final concentration of 0.5 and 1 wt. %. The mixture was frozen overnight at -18 °C and fabricated into a porous structure by the freeze-drying technique. The obtained aerogels were crosslinked in 1 M calcium chloride (CaCl_2_) solution for 1 hour and freeze-dried again to form GOSA scaffolds. The obtained scaffolds are named as GOSA0.5 and GOSA1 according to the concentration of GO in the composite sample. SA scaffolds were fabricated without the addition of GO. In order to make conductive RGOSA scaffolds, the obtained GOSA0.5 and GOSA1 scaffolds were treated with gelatin solution (1 mg mL^-1^) at 95°C to produce conductive RGOSA0.5 and RGOSA1 scaffolds, respectively. The prepared scaffolds were cut in 2 mm thickness (diameter [?]14 mm) until further use. Details on the physicochemical characterization of the GOSA and RGOSA scaffolds can be found in [24].

### 2.3. Scanning Electron Microscopy (SEM)

The cross-sectional area of scaffolds was captured by a scanning electron microscope (SEM, Hitachi, SU1510 high technologies) at an acceleration voltage of 30 kV to observe the morphology and porosity of samples.

### 2.4. *In vitro* biodegradation study

The *in vitro* biodegradation study aims to investigate the biodegradation rate of prepared scaffolds [35]. Briefly, each scaffold (*n* = 3) with known weight (*W*_0_) was incubated at 37degC in alpha-MEM culture media containing 1% (v/v) penicillin/streptomycin to prevent bacterial growth. At specific time points, scaffolds were rinsed with ddH_2_O. Then, samples were dried under vacuum and weighed (*W*_1_). The extent of biodegradation was calculated using the following formula.

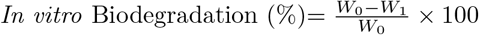

### 2.5. Sterilization and coating of scaffolds

Before cell seeding, the scaffolds were sterilized with 80% ethanol for 24 hours, then allowed to dry overnight and washed twice with DPBS. For coating, scaffolds were incubated at room temperature in poly-l-lysine (PLL) (10 µg/mL) overnight followed by incubation at 37 °C in laminin (LAM) solution (10 µg/mL) overnight. Non-coated (NC) scaffolds, which were wetted but not incubated with any coating material, served as control.

### 2.6. Cell culture

Donor DPSCs (passage 6-7) were cultured in Alpha modification of Eagle’s medium (α-MEM) supplemented with 10% FBS, 1% penicillin (100 U/mL)/streptomycin (100 µg/mL), 100 µM L-ascorbic acid 2-phosphate, and 2 mM L-glutamine at 37 °C in 5% CO_2_ humidified atmosphere. Cells reaching 80–90% confluency were harvested using 0.05% (w/v) trypsin/ethylenediaminetetraacetic acid (EDTA) solution and seeded onto scaffolds in 24-well culture plates (100 µL/well) at 1, 2, 4, 8, or 16×10^4^ cells/sample. The scaffolds were incubated for 3 hours at 37 °C with 5% CO_2_ to allow diffusion of cells through the network of pores before adding culture media (550 µL/well). DPSCs cultured on the surface of the well plate (i.e. No scaffolds) were used as 2D control.

### 2.7. Cell viability using Alamar Blue assay

Alamar Blue (AB) assay was used to quantitively measure the metabolic activity of living cells on the scaffolds by detecting the oxidation-reduction rate of AB reagent [36]. The effects of cell seeding density and coating condition of the various 3D porous scaffolds on the viability of DPSCs were evaluated. Briefly, after 24 and 48 hours of cell seeding on scaffolds, 1 mL of 10% AB solution was added to each well (250 µL/well). The plates were shaken gently (200 rpm) for 5 min and incubated for 4h at 37 °C with 5% CO_2_. After incubation, 100 µL of each sample was transferred to a 96-well plate and the fluorescence intensity was recorded at an excitation wavelength of 540 nm and an emission wavelength of 600 nm using a spectrophotometer (BioTek Synergy H1 multi-mode reader). Non-seeded scaffolds supplemented with 10% AB dye were used as a negative control to confirm that the fluorescence intensity of scaffolds alone did not interfere with the assay. Each experimental condition was conducted in triplicates, and experiments were replicated twice.

### 2.8. Cytotoxicity study using Lactate Dehydrogenase (LDH) assay

The cytotoxic effects and level of toxicity of fabricated 3D scaffolds seeded with DPSCs, considering various coating conditions and inclusion/exclusion of serum in media, were measured by lactate dehydrogenase (LDH) assay for up to two days under proliferation conditions. The LDH kit (CytoTox 96® non-radioactive cytotoxicity assay, Promega) was used and LDH assay was performed according to manufacturer’s instructions in which the number of cytotoxic cells is quantified by measuring cytosolic LDH enzyme leakage into the culture medium as a result of cell membrane damage [37]. Density 3 (4×10^4^ cells/sample) was chosen to compare the DPSCs toxicity of scaffolds. Briefly, 50 µL of cell culture media was collected from 24-well plates after 24 and 48h following DPSC seeding and transferred into a fresh 96-well plate. 50 µL of LDH assay mixture was then added to each well containing the supernatant and incubated at room temperature in dark for 30 minutes. After 30 min, the reaction was stopped with HCl (1 N, 10 vol %) and absorbance values were obtained at 490 nm using a 96-well plate reader (GloMax Discovery microplate reader). DPSCs grown without scaffolds (2D controls) were incubated with lysis solution for 45 min and used as positive controls (100% dead, maximum LDH release control). The cytocompatibility performance of the scaffolds was analysed by using the absorbance of the experimental groups and the negative control group (scaffold only, with no cells). Cytotoxicity data are presented as the average of three replicates. Two different donors were selected to determine the effects of donor type on cytotoxic effects of DPSCs with and without FBS in culture media.

### 2.9. Statistical analysis

Data are graphically reported as mean ± SEM (standard error of mean) of at least three independent samples. Statistical analysis was performed by two-way analysis of variance (ANOVA) with a significance level of*p <* 0.05 followed by post hoc Dunnett test. The analysis was carried out on GraphPad Prism software.

The correlation between cell viability and mean pore size was determined by the Spearman Rank Order Correlation test.

## Results

### 3.1. Structural analysis

The microstructure of fabricated scaffolds was assessed using SEM images (Fig. 1a-e). The data revealed that all scaffolds exhibited homogeneous porous structures with interconnected pores. As reported in our previous study [24], the average pore sizes were measured as 162.5, 147.4, 142.5, 116, and 114.7 µm for SA, GOSA0.5, GOSA1, RGOSA0.5 and RGOSA1, respectively. These observations show the dependency of composite scaffold pore size on GO concentration.

**Fig 1.**
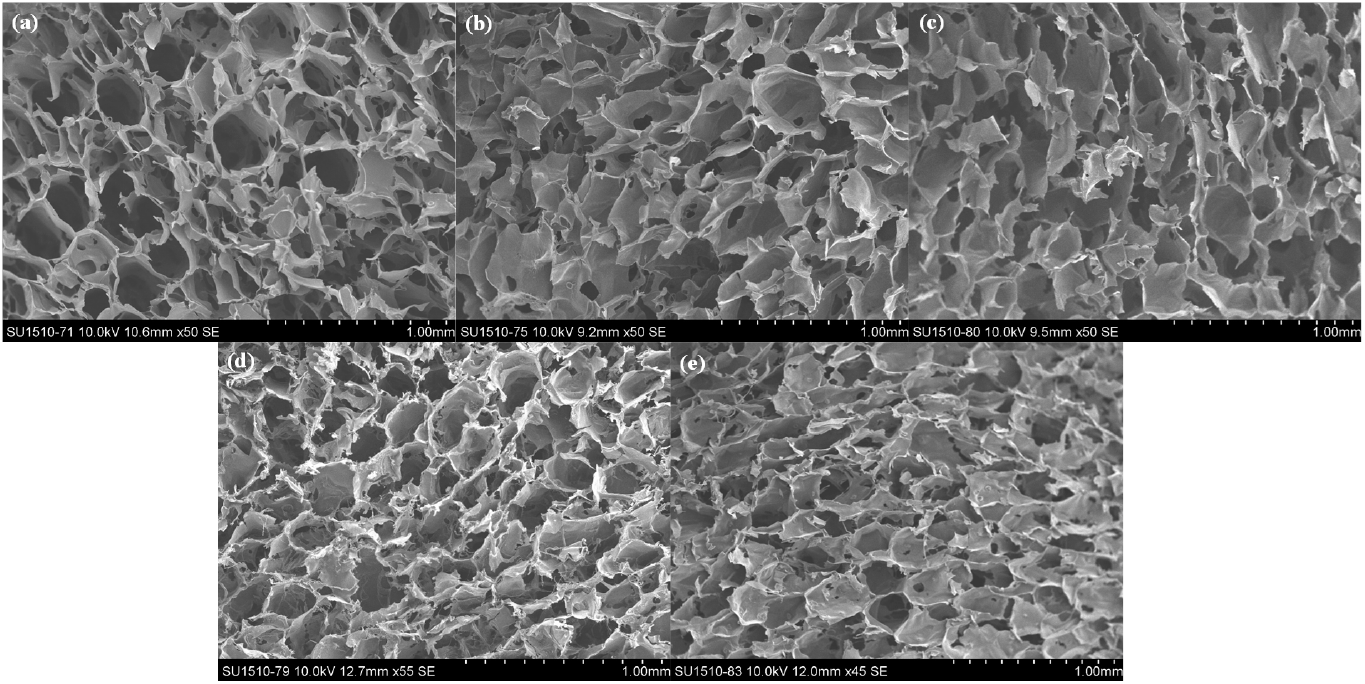
SEM images of (a) SA, (b) GOSA0.5, (c) GOSA1, (d) RGOSA0.5 and (e) RGOSA1 scaffolds showing connected porous structure of fabricated scaffolds to facilitate 3D DPSCs culture.

### Hosted file

`image1.emf` available at https://authorea.com/users/389095/articles/503728-biodegradable-and-biocompatible-graphene-based-scaffolds-for-functional-neural-tissue-engineering-a-strategy-approach-using-dental-pulp-stem-cells-and-biomaterials

### 3.2. Biodegradation study

To evaluate the effect of GO addition on the degradation rate of composite scaffolds, weight loss was expressed as the percentage (%) of biodegradation within three weeks (Fig. 2). The biodegradation of GOSA and RGOSA scaffolds containing different percentages of GO were analysed over a three-week period. At day 3, there was gradual biodegradation of all GOSA and RGOSA scaffolds by approximately 20% of initial weight. In contrast, SA scaffolds showed degradation percentages of approximately 31%. The difference between all GO-based scaffolds and the SA scaffold (no-graphene control) was statistically significant (*p <* 0.0001) from day 2 to day 21. Fig. 2 presents *in vitro* biodegradation data for all five scaffolds. The mean (SEM) weight loss between day-0 and day-21 was approximately 24.86%±1.34, 32.42%±0.68, 23.19%±1.60, and 31.42%±1.20 for GOSA0.5, GOSA1, RGOSA0.5, and RGOSA1 scaffolds, respectively. The SA scaffold (containing no graphene) showed the highest weight loss of approximately 44.02% at day 21, while hybrid scaffold weight loss stabilised after 7 days.

**Fig 2.**
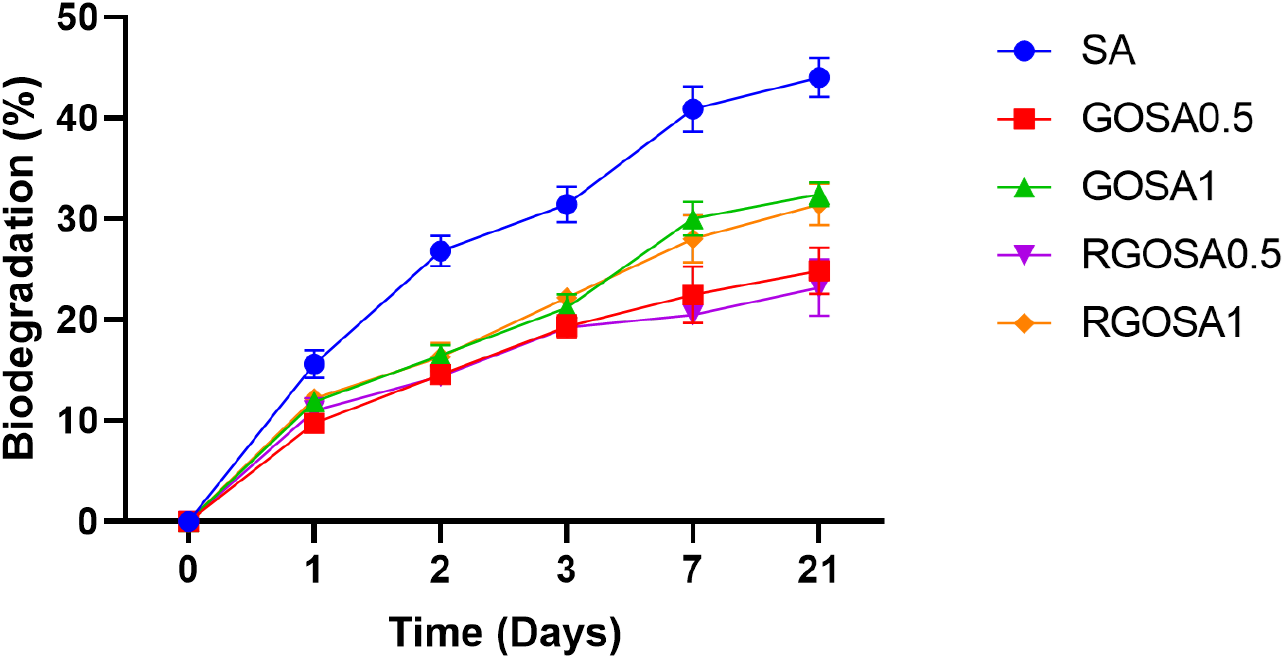
The biodegradation rate expressed as a percentage (SEM) of SA, GOSA0.5, GOSA1, RGOSA0.5 and RGOSA1 scaffolds from 0 to 3 weeks.

#### Hosted file

`image2.emf` available at https://authorea.com/users/389095/articles/503728-biodegradable-and-biocompatible-graphene-based-scaffolds-for-functional-neural-tissue-engineering-a-strategy-approach-using-dental-pulp-stem-cells-and-biomaterials

### 3.3. Enhancement of cell viability using 3D culture system

The viability of DPSCs grown in 2D or 3D scaffolds was assessed using the AB assay. After 24 h of cell culture (Fig. 3), there was a significant increase in the cellular activity of SA and GOSA scaffolds seeded with DPSCs compared to DPSCs grown under 2D conditions at all cell densities (*p <* 0.0001). Cells seeded directly on the surface had an average (SEM) AB reduction of 53.7%±1.00 across all five seeding densities. Furthermore, cells on 3D scaffolds were found to be more viable when compared to the no scaffold (2D) condition (Fig. 3).

**Fig 3.**
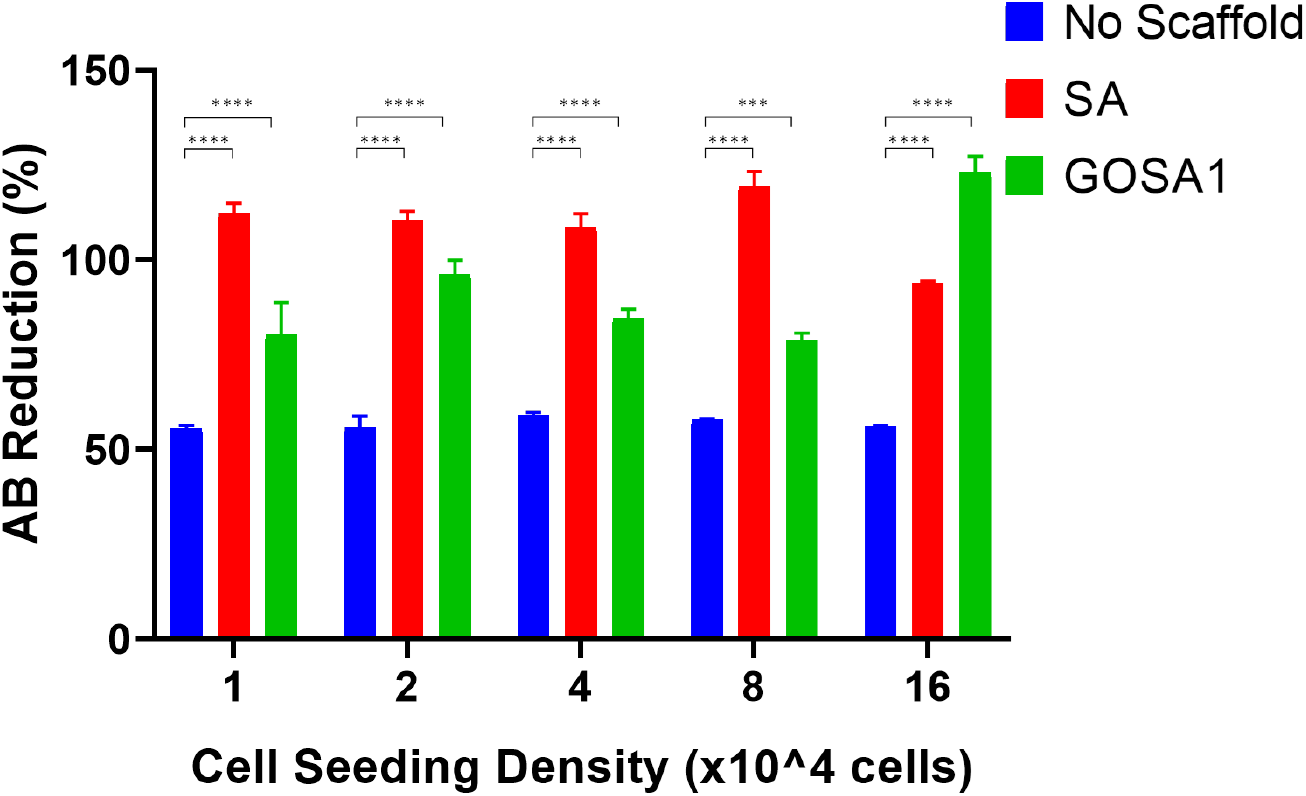
24-hour Alamar Blue reduction (%) of 2D surface (No Scaffold), SA and GOSA scaffolds (means, SEM) at five cell seeding densities; * indicates statistical significance (***p *<* 0.001, ****p *<* 0.0001).

### 3.4. Comparison based on cell densities

Both scaffolds (SA and GOSA) supported cell viability across various cell densities with no negative effect on seeding efficiency, after 24h of cell culture (Fig. 3). The data showed that AB reduction increased significantly, as an indication of metabolic activity, in SA and GOSA1 scaffolds at all five cell densities.

#### Hosted file

`image3.emf` available at https://authorea.com/users/389095/articles/503728-biodegradable-and-biocompatible-graphene-based-scaffolds-for-functional-neural-tissue-engineering-a-strategy-approach-using-dental-pulp-stem-cells-and-biomaterials

### 3.5. Increased DPSCs viability on scaffolds coated with PLL

As DPSCs viability was not affected by seeding density conditions, densities 1, 3 and 5 were selected to conduct AB analysis over 24- and 48-hours. In order to determine the effects of coating conditions on seeded DPSCs, three different coating conditions (NC, PLL, and PLL+LAM) were used. The effect of cell seeding densities was impacted by various coating conditions following 24 hours of culture. Significantly higher AB reduction percentage was observed for SA and GOSA scaffolds at all three densities compared to the 2D control condition (Fig. 4a-c). After 48 h of DPSCs culture, statistically significant differences were detected in the proliferation profiles of cells growing in 2D and 3D under various coating conditions (Fig. 4d-f). In addition, PLL coating significantly increased the cellular activity of GOSA scaffolds at all three cell densities within 48 hours of DPSCs culture (Fig. 4).

**Fig 4.**
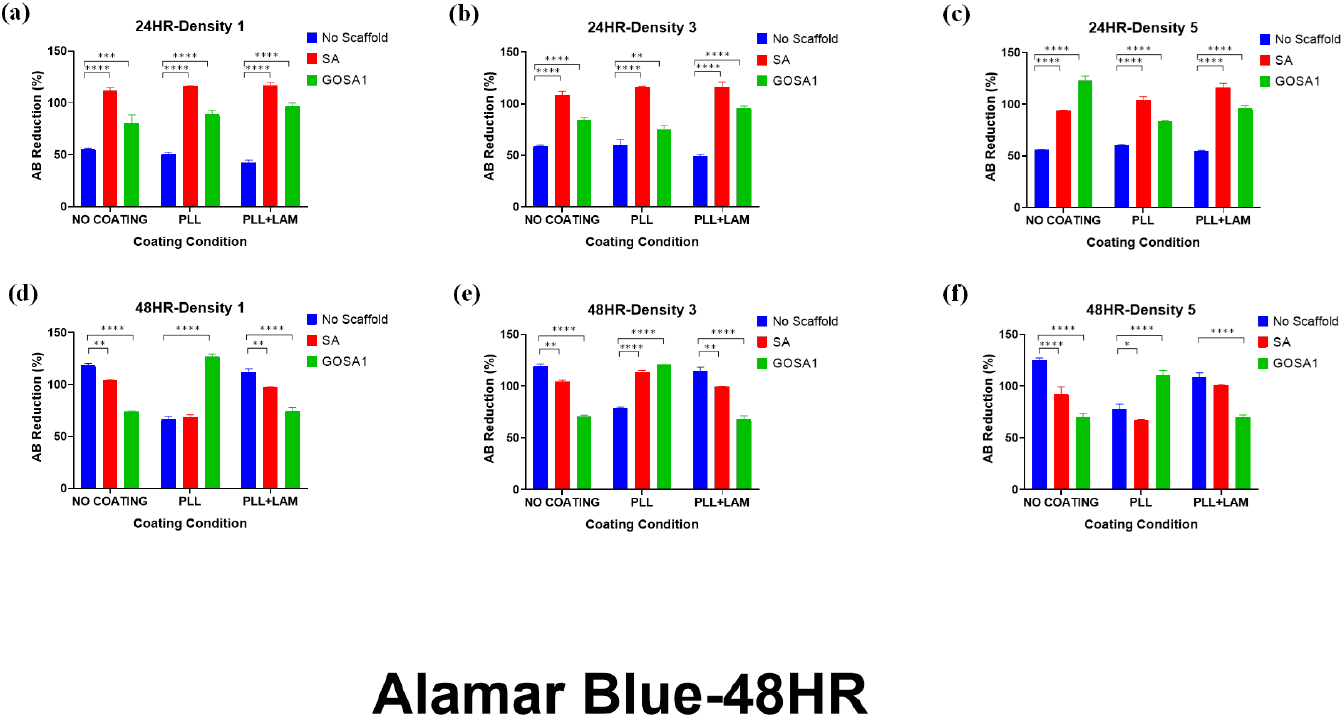
(a-c) 24- and (d-f) 48-hour change (means, SEM) in Alamar Blue reduction percentage of No scaffold, SA and GOSA1 scaffolds across (a, d) Density 1 (1×10^4^cells/scaffold), (b, e) Density 2 (4×10^4^cells/scaffold), and (c, f) Density 5 (16×10^4^cells/scaffold); * indicates statistical significance (**p *<* 0.01, ***p *<*0.001, ****p *<* 0.0001).

#### Hosted file

`image4.emf` available at https://authorea.com/users/389095/articles/503728-biodegradable-and-biocompatible-graphene-based-scaffolds-for-functional-neural-tissue-engineering-a-strategy-approach-using-dental-pulp-stem-cells-and-biomaterials

### 3.6. Scaffold biomaterial composition plays a key role in cellular functions

To explore the effects of scaffold biomaterials on cellular functions, the metabolic activity of cultured DPSCs on various scaffolds was measured using the AB assay. In SA scaffolds, AB reduction (%) was maintained at the 2D cell culture (Fig. 5). In contrast, the AB reduction was significantly increased for GOSA1, RGOSA0.5 and RGOSA1 scaffolds compared to SA only scaffolds across all coating conditions. A significantly higher degree of reduction of the AB dye was observed in DPSCs grown on PLL-coated graphene-based scaffolds compared to those cultured on a 2D surface. This highlights the importance of PLL coating (as previously proved in coating conditions), where the highest AB reduction percentage was recorded for PLL-coated RGOSA1 as 93.66%±5.88.

**Fig 5.**
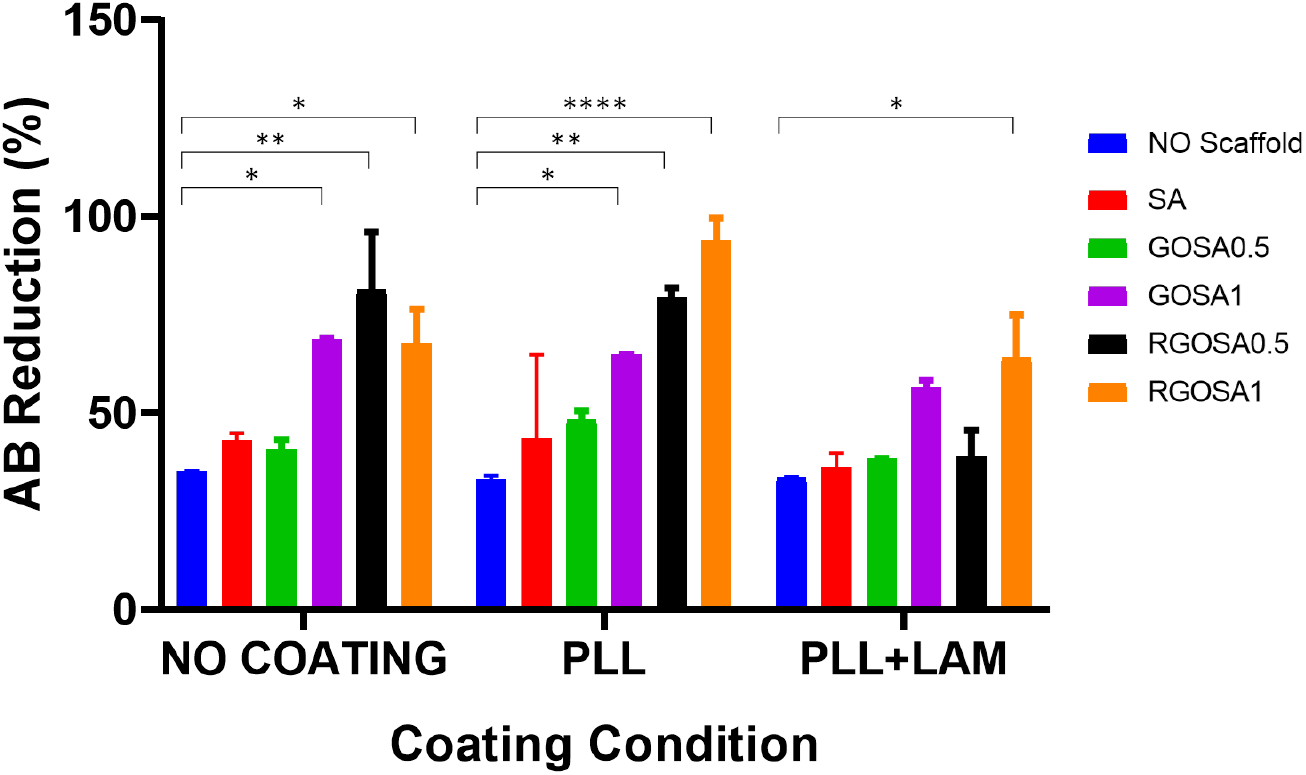
Alamar Blue reduction (%, means, SEM) of DPSCs seeded on 2D surface (No scaffold), SA, GOSA0.5, GOSA1, RGOSA0.5 and RGOSA1 scaffolds after 48h; * indicates statistical significance (*p *<* 0.05, **p *<* 0.01, ****p *<* 0.0001).

#### Hosted file

`image5.emf` available at https://authorea.com/users/389095/articles/503728-biodegradable-and-biocompatible-graphene-based-scaffolds-for-functional-neural-tissue-engineering-a-strategy-approach-using-dental-pulp-stem-cells-and-biomaterials

### 3.7. Lower cytotoxic effects of graphene-based scaffolds

The cytotoxicity of fabricated scaffolds was examined using analysis of lactate dehydrogenase (LDH) release in culture media. In this study, LDH release was taken as a marker of DPSCs membrane damage. At 24 hours of DPSCs culture, all biomaterials showed a relatively low level of released LDH (Fig. 6a). However, significant increase in the LDH levels of DPSCs were observed for uncoated SA (41.65%±7.35) and PLL-coated GOSA0.5 (34.31%±5.31) scaffolds after 24 hours of cell culture. It is interesting to note that, after 48 hours of culture, DPSCs toxicity obtained by LDH assay on graphene-based scaffolds were not significantly higher than the 2D surface (Fig. 6b). However, DPSCs cultivated onto pure SA scaffolds displayed the highest levels of cytotoxicity as compared to 2D control (no scaffold) (*p <* 0.0001).

**Fig 6.**
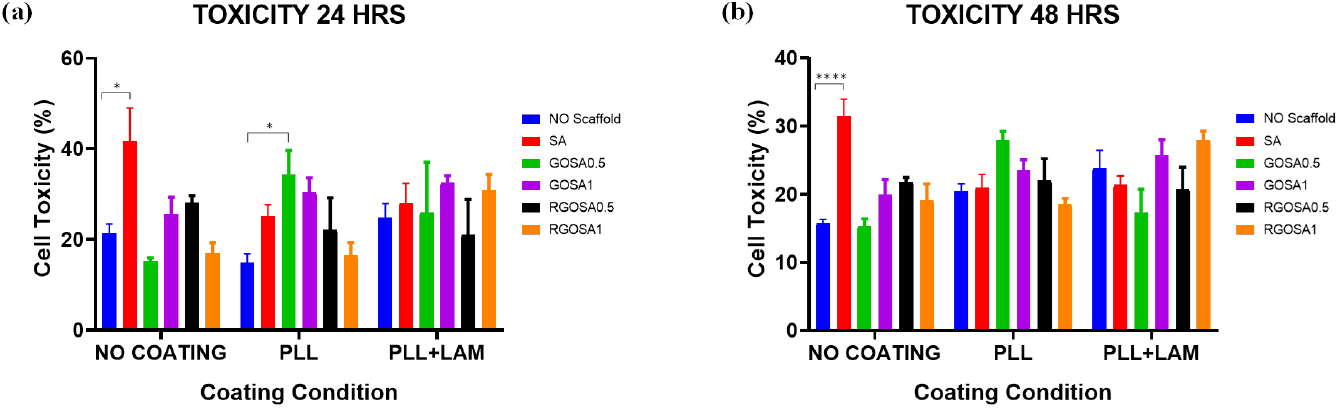
Cell cytotoxicity (mean, SEM) of each scaffold measured by LDH assay in no coating, PLL coating and PLL+LAM coating conditions at (a) 24 and (b) 48 hours of DPSCs culture; * indicates statistical significance (*p *<*0.05, ****p *<*0.0001).

#### Hosted file

`image6.emf` available at https://authorea.com/users/389095/articles/503728-biodegradable-and-biocompatible-graphene-based-scaffolds-for-functional-neural-tissue-engineering-a-strategy-approach-using-dental-pulp-stem-cells-and-biomaterials

### 3.8. Better cellular behaviour in scaffolds with smaller mean pore size

The AB and LDH assay results were utilized to determine the effects of scaffold pore size on cellular behaviour. The data showed that cellular activity decreases when the mean pore size increases on PLL-coated scaffolds (Fig. 7a). The lowest AB reduction percentages were measured for PLL+LAM coated SA and GOSA0.5 scaffolds with the largest mean pore sizes as 36.12% and 38.33%, respectively. Accordingly, the relatively lowest cytotoxicity was observed for RGOSA0.5 (116.0 µm) and RGOSA1 (114.7 µm) scaffolds with smaller mean pore sizes compared to SA and other composite scaffolds (Fig. 7b). Overall, a negative correlation (Spearman R= −0.83) was observed between mean pore size of scaffolds and cellular activity (obtained by AB assay). Conversely, a weak positive correlation (Spearman R= 0.06) was seen between the mean pore size of scaffolds and cytotoxicity (obtained by LDH assay). These findings indicate that increasing mean pore size decreases DPSCs metabolic activity with no corresponding relationship to cytotoxicity.

**Fig 7.**
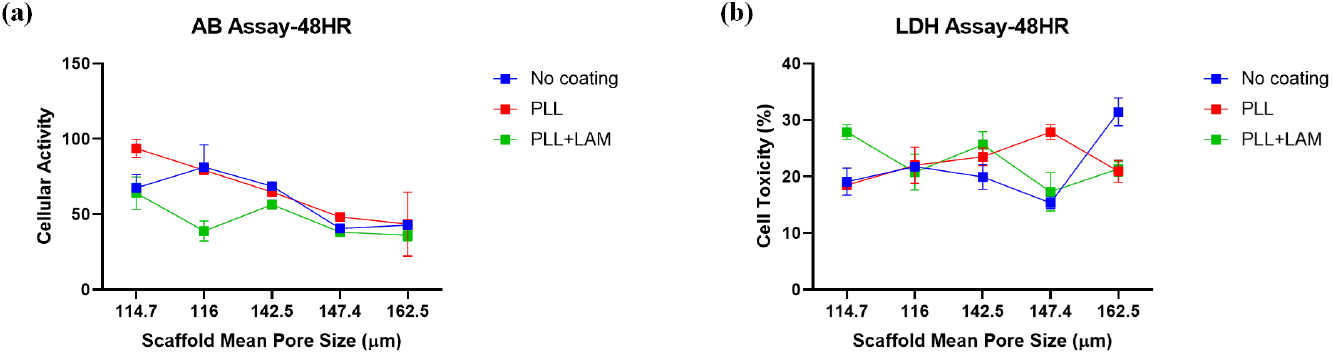
Effect of mean pore sizes on (a) cell activity and (b) cell toxicity (mean, SEM) within various scaffolds (n=3) 48h after incubation (mean pore sizes: SA=162.5 µm, GOSA0.5=147.4 µm, GOSA1=142.5 µm, RGOSA0.5=116.0 µm and RGOSA1=114.7 µm). There is a strong negative correlation between cellular activity and mean pore size (Spearman R = −0.83) and only a weak positive correlation between cell toxicity and mean pore size (Spearman R = 0.06), that is, the enlargement of mean pore size led to a decrease in cellular activity and did not correlate with cellular toxicity.

#### Hosted file

`image7.emf` available at https://authorea.com/users/389095/articles/503728-biodegradable-and-biocompatible-graphene-based-scaffolds-for-functional-neural-tissue-engineering-a-strategy-approach-using-dental-pulp-stem-cells-and-biomaterials

### 3.9. Serum supplemented scaffolds greatly increased cytotoxicity

DPSCs cultured for 24h in serum-free media on all graphene-based scaffolds caused no significant increases in cytotoxicity levels compared to cell only (2D) control, irrespective of coating conditions (Fig. 8a). The data also indicated significantly lower cell toxicity of PLL+LAM-coated RGOSA1 scaffolds (1.81%±3.57), compared to no scaffold condition. Furthermore, quantitative LDH activity measurements (Fig. 8b) showed no significant differences after 24 hours between all 3D graphene-based scaffolds seeded with DPSCs using serum-containing media, when compared to 2D control. These cytotoxic effects were not influenced by coating conditions. However, the percentage of cytotoxic effects of PLL+LAM-coated SA scaffolds (24.83%±4.77) was the highest amongst other scaffolds assessed across all coating conditions.

**Fig 8.**
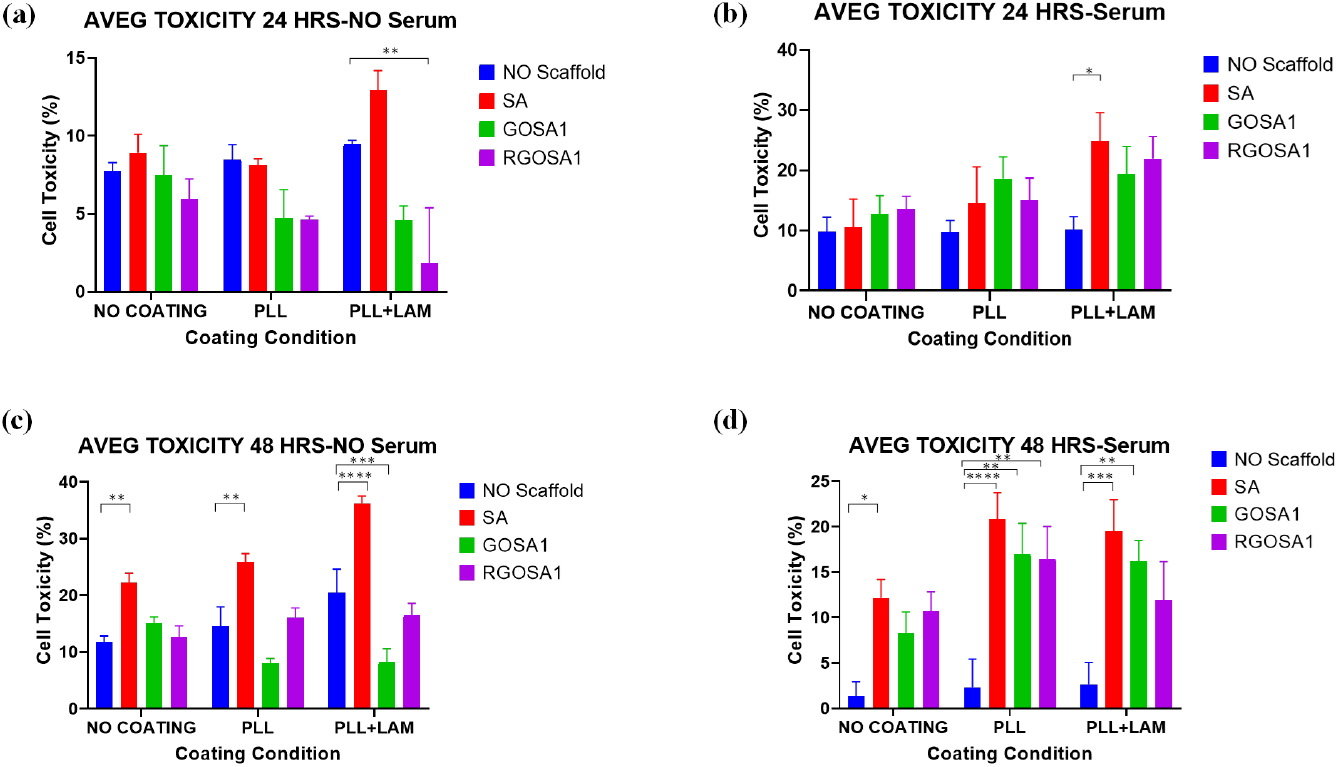
Comparative evaluation of LDH assay (mean of two different donors, SEM) by DPSCs grown on SA, GOSA1 and RGOSA1 scaffolds at (a, c) 24- and (b, d) 48-hours post-seeding in (a, b) serum-free and (c, d) serum-containing media; * indicates statistical significance (*p *<*0.05, **p *<* 0.01, ***p *<* 0.001, ****p *<* 0.0001).

There are no significant increases in the percentage of cell toxicity of DPSCs exposed to coated and uncoated GOSA and RGOSA scaffolds in comparison to the 2D control (Fig. 8c). However, after 48 h of cell culture, a significant elevation of LDH release was detected when DPSCs were cultured on SA scaffolds with serum deprivation, with the highest cell toxicity percentage of 36.22%±1.26 for PLL+LAM coating. The cell toxicity percentage of almost all samples tested with no serum was found to have increased in comparison to the corresponding conditions at the 24 h time point. In addition, when DPSCs in serum-rich media were seeded onto fabricated 3D scaffolds, cell toxicity of scaffolds increased significantly in comparison with no scaffold culture, regardless of coating conditions (Fig. 8d). After 48h of DPSCs culture with serum, cell cytotoxicity was the highest in SA (20.78%±2.95), GOSA1 (16.95%±3.38), and RGOSA1 (16.43%±3.58) matrices coated with PLL. Furthermore, at 48 h of cell culture with serum, the cell toxicity percentage of all evaluated samples was reduced when compared with the corresponding 24 h time point.

#### Hosted file

`image8.emf` available at https://authorea.com/users/389095/articles/503728-biodegradable-and-biocompatible-graphene-based-scaffolds-for-functional-neural-tissue-engineering-a-strategy-approach-using-dental-pulp-stem-cells-and-biomaterials

## Discussion

Significant complexity has been uncovered in the interaction between stem cells and engineered scaffolds [38]. In this study, we report the profiles of DPSCs cultured on fabricated 3D GOSA and RGOSA scaffolds with 2D, coating and media controls.

Our *in vitro* biodegradation data revealed an inverse relationship between GO concentrations and weight loss. This relationship is explained by accessibility of water molecules to GO composites as a function of GO concentration. As discussed in the water contact angle measurements in our previous paper [24], the incorporation of GO increases the interaction of composite scaffolds with water and this effect is GO concentration-dependent. Therefore, as hydrophilicity accelerates with higher GO concentration, water-mediated scaffold degradation increases [39]. Notably, a reduced degradation rate is assumed to be beneficial for tissue regeneration [40]. Collectively, our results confirm that the degradability of composite 3D R/GOSA scaffolds is adjustable and controlled by graphene content.

It is noteworthy that when cultured onto 3D SA and GOSA scaffolds, DPSCs viability was enhanced in relation to 2D culture plates. This relationship clearly signifies the optimal condition of initial cell adhesion to the scaffold surface to promote subsequent cell proliferation and infiltration. Our observation of an increase in the total metabolic activity of cell-seeded scaffolds provides proof-of-principle support for cell growth and proliferation in a 3D matrix which can act as a delivery system for seeded cells. Thus, it is reasonable to conclude that an artificial 3D scaffold is an acceptable approach to mimic the natural architecture of the native tissue and crease a microenvironment conducive to DPSCs engraftment.

The cell-cell interaction within the matrix of 3D cell culture systems has a profound influence on cellular functions including viability, migration and proliferation in contrast to 2D culture [41]. For example, one report showed that 3D polymer-based scaffolds seeded with hepatic cells had less cytotoxic effects than those cultured in 2D [42]. In another study, the 3D culture of dental stem cells was found to support their neuronal characteristics and maintain cell phenotypes [43]. Extending this, we have confirmed that a superior proliferative ability of DPSCs, as measured by metabolic activity, can be obtained when cells are cultured on 3D porous scaffolds.

Regarding cell seeding density, we have shown that when cells are seeded on 3D scaffolds at all densities, the cell proliferation rate is significantly increased in comparison to 2D. Moreover, we found that increased seeding density in 3D scaffolds could be achieved without inducing cytotoxic effects, as determined by LDH assay. In this study, we also showed that the addition of graphene to 3D composite scaffolds improved cellular behaviours and this was seen across all DPSCs seeding densities examined. Notably, the degree of cellular metabolic activity did not differ significantly between the different cell densities tested. This result is important because the implication is that seeding efficiency can be achieved even at high cell densities, at least within a 48-hour period. More work would be required to determine if high seeding densities might have different effects on longer-term culture.

Assessment of coating reagents revealed that PLL+LAM coating did not affect cell viability as indicated by AB reduction percentage. After 48 hours of DPSCs culture, both laminin coating and no coating conditions decreased cell viability on both SA and GOSA scaffolds. Possible explanations involve interactions between coating properties and DPSCs adherence or cell aggregation which can decrease proliferation [44]. Interestingly, PLL was identified as the coating reagent that enhanced cell-matrix adherence. This enhancement might be due to the larger number of cationic sites offered by PLL coating on the 3D surface. In agreement with another study [44], our results show that PLL is superior to laminin coating. In addition, the effect of all three coating conditions on DPSCs was irrespective of cell seeding density.

The AB assessment of metabolic activity shows that biomaterial composition can modulate DPSCs responses to fabricated scaffolds. Our no scaffold controlled LDH results also indicate that SA, GOSA and RGOSA scaffolds materials are nontoxic to DPSCs in short-term culture. Based on previous work, it might be inferred that the composition of a scaffold material can have a direct effect on the biodistribution of secreted factors that in turn influence the stem cell fate. The results appear to suggest that different scaffolds with varying material properties (such as blend ratio, swelling index, or microstructure) elicit diverse DPSCs behaviours. The increase in cell viability observed*in vitro* upon the incorporation of graphene in composite scaffolds is consistent with other studies [36, 45]. Published data suggest that the outstanding surface properties and adsorption capacity of graphene-based nanomaterials are the main contributors to the observed DPSCs responses.

Our results showed a strong influence of pore size, material composition and substrate dimensionality on cell viability, in accordance with the findings of Domingos et al. [46]. Comparisons of AB reduction in SA (97.2%) and graphene-based scaffolds including GOSA0.5 (97.5%), GOSA1 (98.0%), RGOSA0.5 (99.05%), and RGOSA1 (99.18%) revealed the differences in DPSCs proliferation markers across various graphene-based scaffolds. These differences appear to be explained by variations in scaffold porosity (%) such that scaffolds with higher porosity (RGOSA [?] 99%) are able to accommodate higher numbers of viable cells. Furthermore, it was shown that scaffolds with smaller mean pore sizes induce relatively less toxicity. This can be explained by the available surface area of scaffolds for cultured cells or applying the principle that the mean pore size and specific surface area are inversely proportional. In consideration of the specific surface area and mean pore size of a scaffold, biophysical properties can affect cell adhesion [47]. It follows that the low levels of cell adhesion are observed on scaffolds with larger pore size and less specific surface area [48, 49]. As a result, the available specific surface area per unit volume for cell adhesion of each fabricated scaffolds can be calculated using mean pore sizes [50]. Accordingly, the normalized specific surface area of GOSA and RGOSA scaffolds, as shown in Table 1, can be obtained by dividing the mean pore size of each scaffold by mean pore size of SA scaffold (3D control). Thus, the higher AB reduction observed in our RGOSA1 scaffold can be explained by the higher specific surface area in comparison with GOSA1 and SA scaffolds. These data indicate that higher pore size facilitates increased DPSCs migration and proliferation, in agreement with a previous report on culturing DPSCs into 3D poly-L-lactic acid-based scaffolds [51]. In addition, the mechanical properties of scaffolds with overly large pores are compromised, whereas higher cellular proliferation within large pore sizes can have implications for differentiation [52].

**Table 1.**
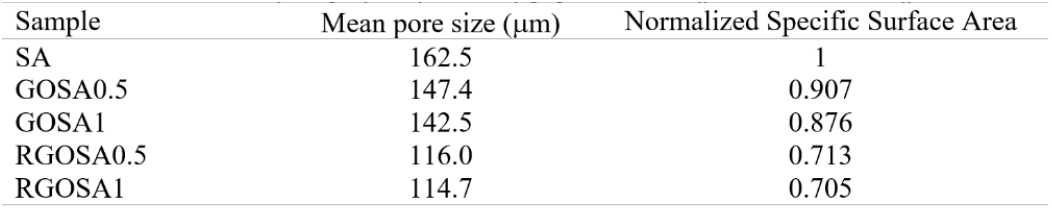
Estimates of the specific surface area of graphene-based scaffolds relative to SA scaffold

Our cytotoxicity findings are consistent with several other studies using GO layer cultured with mesenchymal stromal cells and GO/chitosan scaffolds seeded with human adipose-derived stem cells [53, 54]. In our study, LDH measurements showed significantly higher levels of DPSCs toxicity for uncoated SA scaffolds which can be attributed to poor ability and lack of efficient sites to support cell adhesion and proliferation. This result is supported by another study utilizing other materials mixed with alginate to create biocomposites [55]. However, the incorporation of GO into GOSA scaffolds did not elicit significant differences in cytotoxic effects after 2 days of DPSCs culture. Overall, the GO-enriched scaffolds exhibited cytotoxicity of 15–27% after 48h of culture, suggesting that these materials are biocompatible with DPSCs. Therefore, GO has no apparent cytotoxic effect but exhibits positive effects on cell function in long-term DPSCs culture.

The use of serum-free or serum-rich culture media for biological assays is a contentious issue [56]. The associated clinical uncertainties with the use of FBS include immune rejection, batch-to-batch variations, ethical concerns, interfering effects of unidentified growth factors and proteins, and viral contamination [57]. Regarding DPSCs stability, there is clear evidence about the positive effects of serum-free culture media on retaining stemness [58, 59]. Furthermore, serum-free cultured DPSCs within chitosan scaffolds expressed stem cell markers (Nestin and Sox2) and survived successfully after transplantation into the rodent spinal cord [60]. The present study investigated whether serum depletion facilitates the attachment of DPSCs onto hybrid scaffolds. Thus, the effects of serum on the cytotoxicity of DPSCs were tested using 2D and 3D culture systems across three different coating conditions at a single cell density of 4 × 10^4^cells/sample. We have indicated that the presence of serum caused significantly higher cytotoxic behaviour of coated 3D scaffolds which could be due to suboptimal surface-cell attachment [20]. This was also shown for the combination of PLL and LAM coatings in the presence of serum on 3D structures. It is surmised that coatings lower the available surface area of 3D scaffold biomaterials for the attachment of cell receptors, which subsequently causes poor adhesion. In contrast, coated 2D surface was found to result in better attachment of DPSCs which is due to the unfavourable bare tissue culture surface. Moreover, our study found that the cytotoxicity of DPSCs cultured in serum-rich media on 3D scaffolds increased significantly in comparison with the 2D control. The increased level of cytotoxicity in serum-containing medium could be attributed to the surface oxygen content of GO and RGO, which favours FBS adsorption. This could be due to the formation of protein corona on the GO surface which in turn influence the toxicity of GO-based materials [61, 62]. Furthermore, studies have indicated that serum proteins can have an interfering effect on interactions between nanoparticles, cells and biological molecules [63-65]. Accordingly, Lesniak et al. [64] demonstrated that various protein corona formed on silica nanoparticles modified cell adhesion, cellular uptake, and toxicity, which is determined by the serum protein concentration. It was also shown that serum-containing media resulted in lower cell adhesion and internalization efficiency of silica nanoparticles. The present study found DPSCs can be seeded in serum-free media onto GOSA and RGOSA scaffolds with no cytotoxic effects, showing promising potential for clinical translation as cellular transplants are typically serum-free.

## Conclusion

Alginate-based scaffolds have been extensively investigated for NTE, however, they faced some limitations that hampered their further developments. Importantly, alginate-based scaffolds have poor mechanical strength and degradation rate, which are not matched with native tissue microenvironment. It was found that incorporation of graphene within the alginate matrix can address these drawbacks. However, it is important to investigate cellular viability and cytotoxicity of 3D graphene-based composite scaffold. In addition, it is revealed that coating of scaffolds induces cell functions, adhesion and growth. Therefore, this study examined the biodegradation, biocompatibility, bioactivity and cytotoxicity of neural crest-derived DPSCs loaded graphene-based 3D composite scaffolds, using three different coating conditions.

It was shown that the composite GOSA and RGOSA scaffolds have controlled biodegradability which is effective in therapeutic tissue engineering applications. DPSCs viability cultured onto SA and GOSA scaffolds was higher than that of on 2D controls thus signifying surface cell adhesion followed by cell infiltration through the porous matrices. Therefore, superior proliferative ability of DPSCs can be obtained when cells are cultured on 3D porous scaffolds. The LDH assay showed comparable DPSCs toxicity on the GOSA and RGOSA scaffolds to that obtained on a 2D surface in the absence of the biomaterial, highlighting no significant cytotoxic effects of graphene incorporation after 2 days of DPSCs culture. Furthermore, smaller mean pore size of scaffolds resulted in higher cellular activity and relatively less cytotoxicity, which is due to more available specific surface area on scaffolds with smaller mean pore sizes. In terms of coating conditions, PLL was the most robust reagent that improved cell-matrix adherence and affected metabolism activity of DPSCs, being superior to combined PLL+LAM coating. Furthermore, the cytotoxicity of GOSA and RGOSA scaffolds in the presence of serum is increased compared to serum-free condition, indicating that DPSCs can be cultured in serum deprivation onto the fabricated scaffolds for clinical translation. The findings from the current study suggest that the proposed 3D graphene-based composite scaffolds had a favourable effect on the biological responses of DPSCs which could be exploited in further DPSCs differentiation and electrical stimulation for functional NTE.

## Acknowledgement

This study was supported by AOSpine Asia Pacific Research Grant 2019 (AOSAUNZ(R) 2019-05). The first author wishes to acknowledge the University of Adelaide for awarding Adelaide Scholarship International (ASI) for her PhD study. The authors also appreciate the support from the Australian Research Council Research Hub for Graphene Enabled Industry Transformation (project no. IH 150100003). Also, special thanks go to Mr Arash Mazinani for his support with SEM imaging.

